# NeuroFLAME: A Scalable, Privacy-Preserving Federated Learning Platform for Collaborative, Secure, and Reproducible Multi-Site Neuroimaging

**DOI:** 10.64898/2026.04.30.721881

**Authors:** Dylan Martin, Sandeep Panta, Ross Kelly, Javier Romero, Sunitha Basodi, Bradley Baker, Ravi Teja Girijala, Jessica Turner, Kun Yang, Akira Sawa, Frank Hillary, Anand D. Sarwate, Vince Calhoun

## Abstract

Federated analysis offers a scalable approach to multi-site neuroimaging research by enabling distributed statistical modeling, machine learning, decomposition, harmonization, and validation without exchanging sensitive individual-level data. However, a significant implementation gap exists, as the majority of federated clinical neuroimaging studies remain technical proofs of concept, struggling to fully navigate the rigorous data-sharing regulations and security constraints of real-world clinical and research environments. Compounding this, existing federated analysis frameworks are typically exposed as command-line tools and configuration files, imposing a considerable setup burden that demands specialized technical expertise from neuroscience researchers. To address these barriers, we present NeuroFLAME, an enterprise-grade, open-source federated neuroimaging platform built upon the NVIDIA FLARE (NVFlare) framework. NeuroFLAME couples a client-outbound-only communication framework with a graphical user interface specifically tailored for neuroscientists. Using certificate-based trust and containerized execution, the platform ensures reproducible analyses with baseline privacy guarantees through data localization and mTLS encryption; advanced protections such as differential privacy and homomorphic encryption are available as optional configurations. By ensuring that clients only need outbound communication and that raw data never leaves each site, NeuroFLAME is designed to support institutional data governance requirements and regulatory frameworks such as HIPAA and GDPR. We demonstrate the platform’s utility through three federated analysis workflows, each implemented as a NeuroFLAME computation: multi-site federated closed form voxel-wise regression, decentralized constrained joint independent component analysis (dcjICA) and federated label-based dimensional prediction. Empirical validation of both workflows shows that NeuroFLAME achieves high consistency with established centralized approaches, effectively bridging the gap between experimental federated learning prototypes and production-ready collaborative tools for privacy-preserving, large-scale federated neuroimaging analysis.

## 1 Introduction

Multi-site neuroimaging studies are increasingly essential for detecting subtle brain-based effects, improving statistical power, capturing demographic and clinical heterogeneity, and validating biomarkers across institutions [1, 2]. However, centralizing individual-level imaging and clinical data can conflict with institutional governance policies, data-use agreements, and privacy regulations regarding protected health information such as HIPAA and GDPR [1, 3]. Federated approaches address this challenge by allowing sites to keep data locally while contributing to a shared computational objective through the exchange of approved derivative outputs rather than raw data [4–6].

We use **federated analysis** as the umbrella term for distributed analytic workflows in which individual-level data remain at local institutions and only approved derivative outputs are exchanged. These outputs may include summary statistics, sufficient statistics, model parameters, gradients, component estimates, centroids, harmonization parameters, quality-control metrics, or derived features. Under this broad definition, federated analysis includes both one-pass or low-communication aggregation and multi-round iterative optimization, spanning statistical modeling, harmonization, decomposition, prediction, validation, and workflows often described as federated learning. We use **federated learning** when referring to the broader literature and to computations focused on shared model fitting or representation learning. This distinction is operational rather than categorical, since regression, independent component analysis, dimensional prediction, and neural-network training can all involve models learned from data and may be implemented within a federated analysis framework.

To bridge the gap between federated learning research prototypes and production-ready tools for multi-institution clinical research, we present NeuroFLAME, a graphical user interface (GUI)-based federated neuroimaging analysis platform designed to operate under real-world clinical Information Technology (IT) and governance constraints. Throughout the manuscript, we use computation to refer to a containerized decentralized algorithm that can be deployed and executed across participating NeuroFLAME sites. Built on NVIDIA FLARE (NVFlare) [7], NeuroFLAME emphasizes certificate-based trust, auditable computation flows, and outbound-only communication at client sites to align with hospital firewall constraints. The platform couples this secure substrate with a neuroscience-specific orchestration layer, providing a user-friendly interface that lowers the barrier to entry for clinical and neuroscience researchers. This work makes four primary contributions:

- **The NeuroFLAME Platform**: A modular, GUI-based federated neuroimaging analysis platform designed to operate under real-world clinical IT and governance constraints.
- **A Secure Computation Model**: A reproducible, containerized workflow model built on top of NVFlare that enables secure, outbound-only federated analysis.
- **Standardized Local Pre-processing**: An offline, containerized VBM preprocessing tool that lets independent sites prepare structural MRI data through an identical pipeline without network access, establishing the cross-site comparability that federated analysis presupposes.
- **Empirical Validation**: Validation demonstrated via derated closed form voxelwise regression, decentralized constrained joint ICA [8] and federated Label-Based Dimensional Prediction (Fed-LAMP) [9], demonstrating feasibility and consistency with established centralized approaches.

## 2 Motivation

Multi-site collaboration is increasingly critical in clinical neuroimaging research, where larger and more demographically diverse datasets improve the detection of subtle biomarkers, increase statistical power for psychiatric and neurological conditions, and ultimately accelerate the translation of findings into clinical practice [1]. Despite growing enthusiasm for federated learning (FL) in neuroimaging and clinical research, the transition from research prototype to clinical deployment remains persistently slow. A comprehensive review of FL implementation challenges conducted through May 2024 found that the vast majority of published FL systems are not yet appropriate for clinical use, citing methodological flaws, non-Independent and Identically Distributed (non-IID) data heterogeneity across institutions, communication overhead, and a lack of standardization across implementations [10].

A systematic review of FL applications across healthcare [11] demonstrates both the field’s profound potential and its current limitations. Of the 22,693 articles screened up to August 31, 2023, only 32 studies reported real-life clinical applications. The overwhelming majority (approximately 94.8%) remain technical proofs of concept, often using simulated federated environments rather than real-world multi-institution collaborations. This gap persists through the most recent literature: a 2025 review of FL in public health found that of 4,720 deduplicated records spanning January 2020 through June 2025, only 19 studies qualified as real-world deployments [12], and a 2026 neuroimaging-specific study explicitly notes that FL is rarely deployed in real-world brain imaging scenarios despite its clear potential [13]. Despite this, the field is expanding rapidly across all domains — FL publications grew more than 8-fold between 2020 and 2023, from 722 to nearly 6,000 papers [14] — a trend that is reflected in, but not yet matched by, real-world neuroimaging deployments.

The gap between potential and practice is especially pronounced in neuroimaging data analysis. Institutions routinely collect large, complementary datasets, such as multi-site functional Magnetic Resonance Imaging (fMRI) cohorts, longitudinal structural MRI studies, richly phenotyped clinical registries, yet these remain siloed because the technical and regulatory burden of aggregating them is prohibitive [1]. A single-site neuroimaging study is almost always statistically underpowered for the detection of subtle effects in complex psychiatric and neurological conditions, while centralizing data across sites runs directly into data sovereignty, HIPAA/GDPR restrictions, and institutional review constraints [1, 2]. Federated analysis offers a principled solution: each site retains local control of its data while contributing to a shared statistical estimate, model, decomposition, harmonization procedure, or validation workflow. However, realizing this in practice requires platforms that can navigate real hospital IT environments not just simulated multi-client scenarios run on a single server.

As detailed by [5], FL allows hospitals to collaboratively train models for disease diagnosis, such as tumor segmentation in MRI and Computed Tomography (CT) scans or COVID-19 detection, by sharing model parameters rather than sensitive electronic health records (EHRs). This decentralized approach supports privacy and compliance with regulations such as the HIPAA, while improving model generalizability by using diverse data from varied demographics and enabling cross-institutional collaboration. Recent literature [2] underscores a “huge potential” for FL in mental health research, yet emphasizes a critical need for greater efficiency and accuracy in real-world settings. A persistent challenge is the heterogeneity of data types encountered across institutions - neuroimaging, EHRs, genomics, and histology which necessitates flexible and adaptive analytical pipelines tailored to the scientific objectives of each study.

Existing general-purpose FL libraries provide algorithmic primitives but require significant engineering effort to configure for clinical research environments. Neuroscience researchers are often hindered by a steep learning curve when adapting command-line tools and libraries designed for general use [11]. Researchers have also noted that the complexity of current federated learning tools creates usability barriers for non-technical domain experts, underscoring the need for more accessible, GUI-based interfaces [15].

## 3 Related Work

The field of FL in neuroimaging has evolved through several platforms, each addressing different aspects of data sharing, privacy, and computational accessibility. Command-line-based frameworks such as Fed-BioMed[16], PySyft [17], FedML [18], and MetisFL [19] provide robust algorithmic primitives for federated learning and distributed model execution [4, 20, 21]. However, these tools often require significant programming expertise and manual configuration to navigate the specific security and data-handling requirements of clinical research. While these frameworks are well-suited to users with strong software development backgrounds, they may be less accessible to researchers without substantial programming experience. In this work, we focus specifically on neuroimaging-oriented federated analysis tools that utilize GUI platforms to lower the barrier to entry for domain experts.

Brainlife [22] is a cloud-based open-source platform for neuroimaging analysis, data management, and reproducible workflows. It supports MRI, electroencephalogram (EEG), magnetoencephalography(MEG) and other modalities, and offers full provenance tracking of data and derivatives. Its strengths include ease of use, a broad and reusable workflow library, scalability via cloud or high performance computing (HPC) resources, and support for standard data formats and reproducible research practices. However, because data must be uploaded to the cloud, Brainlife does not satisfy the “data remains on-premises” constraint often required in clinical or tightly regulated institutional environments, making it unsuitable for privacy-preserving federated analyses.

Neurobagel [23] provides a complementary, metadata-focused solution that enables decentralized dataset harmonization and subject-level cohort search across distributed neuroimaging datasets without requiring raw data to be centrally aggregated. Neurobagel uses semantic web standards (e.g. Brain Imaging Data Structure(BIDS), Neuroimaging Data Model(NIDM)) and a modular tool stack to allow institutions to expose harmonized metadata while keeping their data locally. However, Neurobagel is not an analysis or computation platform and does not execute federated computations itself, and therefore does not replace platforms built for decentralized or federated analysis.

COINSTAC (Collaborative Informatics and Neuroimaging Suite Toolkit for Anonymous Computation) pioneered decentralized neuroimaging workflows by introducing a GUI that enables researchers to perform multi-site analyses while keeping source data local [24, 25]. It has demonstrated significant utility in large-scale collaborative projects, such as those within the ENIGMA consortium [26], by producing statistically comparable results to centralized analyses without raw data exchange. Similarly, large-scale collaborative projects involving federated Voxel-Based Morphometry (VBM) analysis across 7 sites with schizophrenia data have demonstrated the utility of COINSTAC [27].

Drawing from our experience with COINSTAC, we observed that neuroscience researchers benefit from tools that abstract away the technical complexity of federated infrastructure while preserving transparency about the statistical and computational workflow. NeuroFLAME aims to bridge this gap by pairing NVFlare’s [7] enterprise-grade infrastructure with a neuroimaging-focused GUI, realizing the benefits of privacy-preserving, large-scale collaborative research [28].

## 4 NeuroFLAME Platform – Architecture

NeuroFLAME is designed for environments where enterprise governance and clinical IT policies are paramount. Built on NVFlare, it emphasizes certificate-based trust, auditable computation flows, standardized versioned modules, and outbound-only communication at client sites. These features are specifically aligned with institutional firewall constraints and compliance expectations. The platform couples this secure substrate with a neuroscience-specific orchestration layer, including a central API (Application Programming Interface) and a desktop application for managing studies and deployments across institutions.

Our primary contribution is a graphical interface through which researchers interact with this infrastructure, organized around how collaborative neuroimaging studies are run by most principal investigators: they define projects and invite sites, configure analysis parameters, select from commonly used neuroimaging algorithms, and inspect results, without writing code or editing configuration files. This workflow is described in detail in Section 4.8.

### 4.1 System Design and Components

The architecture is organized around a modular design that separates orchestration, communication, and computation into distinct layers (Figure 1). The platform is divided into two primary domains: Central Services and Researcher-Side Components.

**Fig. 1:**
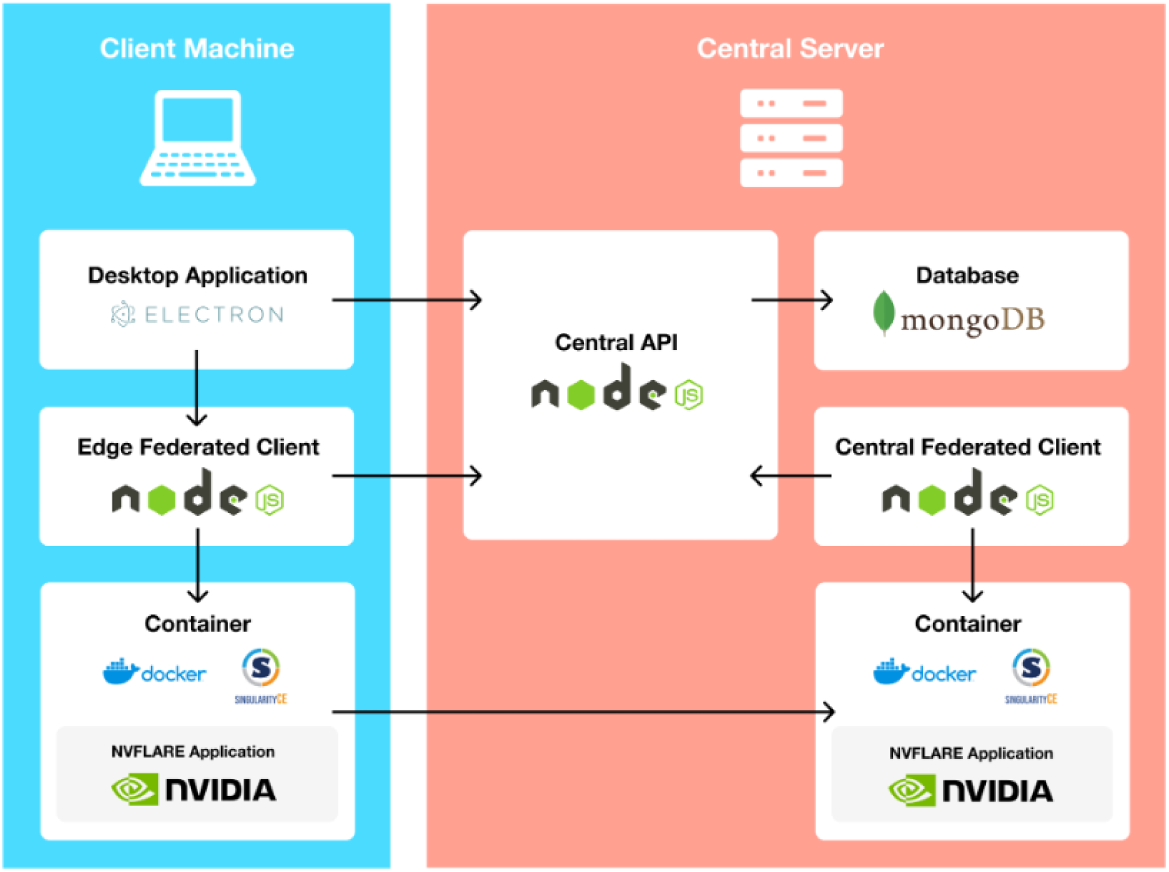
NeuroFLAME architecture showing the relationship between Central Services and local Client Machines.

#### 4.1.1 Central Services

The central infrastructure manages the global state and coordination of the federated network.

- **Central API**: Built with Node.js, this component coordinates study setup, manages consortium membership, and orchestrates active runs.
- **Central Federated Client**: Supervises distributed execution and synchronizes participants across the network. The central server has one inbound port — handling both FL client and admin console traffic.
- **Database and Storage**: A MongoDB instance stores study metadata, while a File Server distributes “runkits” containing configurations and dependencies needed for computation.

#### 4.1.2 Researcher-Side Components

These components operate locally on consortium machines to ensure raw data remains on-premises.

- **Desktop Application**: An Electron-based [29] interface provides a user-friendly environment for joining consortia, configuring study parameters, and monitoring active runs.
- **Edge Federated Client**: A Node.js-based [30] client responsible for pulling manifests and launching the containers that encapsulate the computational workload.
- **Containerization Layer**: Uses Docker [31] or Singularity [32] to instantiate the NVFlare environment, ensuring a reproducible and isolated execution space for the analysis.

#### 4.1.3 Implementation Stack

The platform is implemented with a modern, open-source stack:

- **Frontend and Orchestration**: The desktop application, federated clients, and Central API are written in **TypeScript**. The API exposes **GraphQL** [33] queries and mutations backed by a **MongoDB** [34] database, while the user interface is built in **React** [35].
- **Containerization**: NeuroFLAME employs **Docker** [31] containers (with **Singularity** [32] compatibility for HPC environments) to ensure consistent environments across sites.
- **Analysis**: Inside these containers, federated computations are authored in **Python** [36], the dominant language for machine learning and neuroimaging, though other programming languages are also supported.

### 4.2 Vaults: Always-On Edge Federated Client

A NeuroFLAME Vault is an edge federated client, always online with a curated dataset and ready to participate in federated analyses. Architecturally, a Vault is indistinguishable from any other NeuroFLAME edge client: it holds a local dataset, runs computations inside the same NVFlare containerized environment as all other participating sites, and transmits only model parameters or summary statistics (never raw data) to the central aggregator. Like all edge clients, each Vault is provisioned with its own consortium-specific public key infrastructure (PKI) Startup Kit and mTLS(Mutual Transport Layer Security) credentials [37]. Because Vaults operate as standard edge federated clients, their threat model and security guarantees are fully inherited from the NVFlare architecture described in detail in Section 4.6.

A primary motivation for the Vaults feature is to support scenarios in which datasets cannot be centralized due to regulatory, institutional, or contractual constraints such as Institutional Review Board (IRB) restrictions, Data Use Agreements (DUAs), or institutional governance policies. In these settings, researchers may be authorized to execute approved computational workflows within a controlled environment, but are not permitted to transfer or pool raw data across sites. Vaults enable such researchers to contribute their data to federated analyses while remaining fully compliant with these constraints. A secondary benefit of the always-on design is that it removes the coordination burden of requiring all participating institutions to be simultaneously available; single-site researchers can participate in federated analyses and integrate Vault data with their own local datasets to significantly enhance statistical power and sample size.

Vaults provide access to curated pre-processed neuroimaging datasets along with metadata, ready for federated analysis. Researchers can run computations on Vault data independently or combine it with their own local data within the same federated run.

#### 4.2.1 Hosting and Deploying Vaults

Vaults may be hosted either on secure cloud infrastructure or on-premises at the data-holding institution. Vault owners retain full control over access permissions, configuring their vault as openly accessible to all registered NeuroFLAME users or restricting access to designated collaborators. The vaults currently available through NeuroFLAME are hosted on secure Amazon Web Services (AWS) [38] cloud infrastructure managed by our research center, with datasets pre-processed, quality-controlled, and configured to run specific federated computations as decided by the vault owner. To deploy a dataset as a NeuroFLAME Vault, the vault owner must install the NeuroFLAME Vault client and Docker [31] or Singularity [32] on the hosting machine, then set up the configuration and credentials for their dataset. Once deployed, the vault appears within the NeuroFLAME interface to any users the vault owner has granted permission to, and is listed as an available participant when setting up a federated run. The vault owner can revoke or modify access at any time.

### 4.3 Standardized Local Pre-processing

A recurring obstacle in multi-site neuroimaging is that data must be comparably pre-processed before federated analysis is meaningful. Differences in segmentation software, normalization templates, smoothing kernels, or quality-control criteria introduce site effects that are difficult to distinguish from biological effects in a pooled model [39]. Centralized studies address this by processing all images through a single pipeline at a single location. Post hoc harmonization methods such as ComBat [40] can remove residual site effects from derived features, but they cannot recover comparability that was lost through divergent pre-processing of the images themselves.

To address this, we distribute the NeuroFLAME Preprocessor [41], a containerized voxel-based morphometry (VBM) pipeline for structural MRI. The tool packages SPM12 Standalone [42] together with the MATLAB Compiler Runtime (MCR) [43] inside a Docker [31] and Singularity [32] images, so that no MATLAB licence is required at participating sites and the executed pipeline is identical across institutions. Given T1-weighted images, it performs tissue segmentation into gray matter, white matter, and cerebrospinal fluid; spatial normalization to the standard Montreal Neurological Institute (MNI) template; smoothing; and correlation-based quality control, producing analysis-ready gray matter maps together with the metadata required by NeuroFLAME computations.

#### Offline operation

The preprocessor executes entirely locally and requires no network connectivity at any stage. It does not transmit any data, contacts no external service, performs no licence check or telemetry call, and can be run to completion on a isolated node inside a hospital or institutional network. This is a deliberate design constraint rather than an incidental property. Sites whose governance policies prohibit outbound connections from machines holding protected health information can therefore prepare their data without seeking a network exception. It also draws a clean boundary around the point at which any information leaves a site: pre-processing is a purely local operation, and only the approved derivative outputs of the subsequent federated computation are ever exchanged (Section 4.6).

#### Invocation and shared parameters

The preprocessor is a standalone command-line tool, invoked independently of the NeuroFLAME desktop application and requiring no NeuroFLAME account, consortium membership, or active study. Use of the preprocessor is optional. Sites run it on their own schedule, before or independently of any federated analysis, and may use it purely locally if they never intend to federate the resulting data. This feature also suits Vault owners and sites processing large cohorts, for whom batch invocation and submission to high-performance computing schedulers are more practical than an interactive interface. The tool imposes no requirements on a site’s data beyond an expected input layout, documented with worked examples in the repository [44]; any dataset conforming to that layout can be processed without further configuration.

Because the tool is run independently at each site, cross-site comparability rests on a convention rather than on enforcement: consortium members agree on a common set of pre-processing parameters in advance and each apply that same configuration when invoking the tool. This mirrors established practice in multi-site collaborations, where principal investigators (PIs) agree on the parameters that must be shared for group analysis to be valid, such as the segmentation approach, the normalization template, and the smoothing kernel, while retaining local control over parameters that are necessarily dependent on the properties of each site’s own data, such as acquisition-specific settings and site-level quality-control criteria. Separating the two allows a consortium to commit to a single agreed pipeline without requiring every site to pretend its data were acquired identically. The parameters are supplied to the container as a single configuration file, so that a study lead can circulate one file to all participating sites and each site can verify that the shared settings it applied match those of its collaborators. In the analysis reported in Section 6.1, all three datasets were prepared this way; the agreed protocol is described in Section 6.1.2. We note that this places responsibility for consistency on the consortium, and that the platform does not currently verify that sites in fact used matching parameters.

#### Future development

Exposing pre-processing within the NeuroFLAME desktop application as a dedicated pre-processing computation is a defined priority for future development. Because the Edge Federated Client already retrieves manifests and launches containerized workloads locally (Section 4.1), the same mechanism can distribute a pre-processing configuration specified once by the study lead and execute it at each site, propagating a single agreed parameter set for the pipeline through the interface rather than by circulated file. This would particularly suit consortia whose members want to run same pre-processing pipeline over their data without configuring it themselves. The standalone command-line tool will continue to be supported for batch and highperformance computing use, for sites that prefer to prepare their data entirely outside the platform, and for consortia that require the offline guarantee described above.

We also intend to extend the preprocessor beyond structural MRI to the other modalities routinely combined in multi-site studies, including functional magnetic resonance imaging (fMRI), both task-based and resting-state, and diffusion tensor imaging (DTI).

### 4.4 Integrating NVFlare

A core design decision in NeuroFLAME is the delegation of federated communication and orchestration to NVFlare [7], NVIDIA’s open-source software development kit (SDK) for federated learning and distributed model execution. NVFlare provides secure communication protocols, orchestration tools, privacy-preserving extensions, and development utilities that NeuroFLAME uses to support federatd neuroimaging analysis. It is an enterprise-grade, domain-agnostic SDK that provides a secure, scalable foundation for federated learning and related distributed analytic workflows, including built-in support for advanced privacy-preserving methods like differential privacy and homomorphic encryption. By building upon this infrastructure, NeuroFLAME addresses the need for production-ready collaborative tools that can navigate clinical IT constraints, such as outbound-only communication at client sites and certificate-based identity management. This integration allows the platform to focus on neuroimaging-specific orchestration while inheriting the reliability and security of a proven federated learning backbone [45].

### 4.5 Key Benefits

The architecture of NeuroFLAME is specifically designed to overcome the logistical and technical hurdles that frequently stall multi-institutional neuroimaging collaborations:

- **Privacy-Preserving Design**: NeuroFLAME ensures that raw data remain local, while only approved outputs such as summary statistics, sufficient statistics, model parameters, gradients, centroids, component estimates, harmonization parameters, quality-control metrics, or derived features are exchanged according to the specific computation.
- **User-Friendly Interface**: The neuroscience-focused GUI encapsulates complex implementation details, lowering the barrier to entry for neuroscientists and domain experts who may not have extensive command-line or DevOps (software development and IT operations) expertise.
- **Reproducibility**: Containerized computation (Docker [31] or Singularity [32]) ensures that analysis pipelines are executed in identical, deterministic environments across all participating institutions.
- **Enterprise-Grade Security**: The system utilizes certificate-based identity management and outbound-only mTLS communication at client sites (the central server requires a single inbound port), allowing it to function within highly restricted clinical IT environments and hospital firewalls. The central server requires a single inbound port for federated coordination. NVFlare’s security architecture addresses identity security, communication security, message serialization, data privacy protection, and auditing as five distinct and independently configurable layers [46].
- **Scalability and Performance**: By leveraging the NVFlare backbone, the platform is capable of orchestrating large-scale studies across numerous sites without significant performance degradation.
- **Simplified Development and Standardization**: The platform distributes a containerized local pre-processing tool (Section 4.3) that allows every participating site to prepare its data through an identical pipeline without transferring data or requiring network access, and provides boilerplate templates to facilitate the contribution of new federated methods from the scientific community.
- **Open Source**: All code is available via GitHub at https://github.com/NeuroFlame/NeuroFLAME.

### 4.6 Threat Model and Security Guarantees

NeuroFLAME’s security model is inherited from NVFlare’s security framework [46]. Raw data never leaves local sites; instead, only approved derivative outputs, such as model parameters, aggregated updates, summary statistics, or component estimates, are exchanged between clients and server during federated analysis. This constitutes *privacy-by-architecture*: sensitive data is decentralized by construction, not by policy alone. NVFlare’s framework assumes physical security is already in place for all participating server and client machines, and that mutual TLS provides the authentication mechanism within that trusted environment [46]. Security concerns outside this boundary including physical security, firewall policies, and data storage and retention policies are explicitly delegated to each site’s own IT infrastructure [46].

NVFlare does not inherently protect against stronger adversaries, including: (a) *gradient inversion attacks*, in which a compromised server attempts to reconstruct training samples from transmitted model updates [47]; (b) *membership inference attacks*, in which an observer attempts to determine whether a specific individual’s data was used in training; and (c) *colluding clients*, who might collectively pool the messages they receive to infer site-specific data. These attack classes remain active research areas and are not fully mitigated by the baseline NVFlare configuration [47].

Within this trust boundary, NVFlare implements security across six areas, four of which are active by default and two of which require explicit configuration.

#### Identity and Authentication (active by default)

NVFlare uses a Public Key Infrastructure (PKI) model. For each federated computation, a Project Administrator uses NVFlare’s provisioning tool to generate a root Certificate Authority (CA), which then issues X.509-compliant certificates (2048-bit private keys, 360-day expiry) for every participant, including the FL server, each client site, and administrative users [37]. Each participant receives a separate password-protected Startup Kit containing credentials for mutual TLS (mTLS) authentication, with every file signed by the root CA. In the NeuroFLAME context, Vault servers (Section 4.2) and client sites are each provisioned with consortium-specific credentials, so that no participant can connect without a valid Startup Kit issued for that consortium. Once the federated analysis workflow begins, all inter-node communication is secured via mutually-authenticated TLS (mTLS), so no message can be sent or received without a valid, CA-signed certificate a[37]. NVFlare additionally supports integration with institution-specific authentication systems such as OAuth and Keycloak via an event-based plugin framework, enabling each site to enforce its own identity requirements without modifying the central server [37].

Governance boundaries are enforced through NVFlare’s federated authorization system: every administrative command is subject to per-site authorization policies, and the server is not required to maintain a centralized user list but authorization is evaluated dynamically at each site [37]. This design allows institutions with distinct IT governance frameworks to participate in the same study without ceding local policy control.

#### Communication Security via mTLS (active by default)

Mutual TLS is NVFlare’s default connection mode: the server and clients authenticate each other during connection setup, ensuring that only credentialed clients can connect and that all intermediate statistics and model parameters transmitted during analysis are encrypted in transit [46]. For institutional environments where mTLS is constrained by IT policy, NVFlare supports one-way TLS or Bring Your Own Connectivity (BYOConn), allowing integration of custom networking solutions provided they maintain confidentiality, integrity, and explicit message authentication.

#### Message Serialization via FOBS (active by default)

NVFlare’s Flare Object Serializer (FOBS) is active by default in all current NVFlare versions, replacing the Python pickle format used in earlier releases and eliminating an associated class of deserialization vulnerabilities [46].

#### Auditing (active by default)

NVFlare implements a structured audit logging system that records events at four distinct locations for every job [48]: a *Server Parent* log capturing system-level server events; a *Server Job* log capturing job-specific server events; a *Client Parent* log capturing system-level events at each client site; and a *Client Job* log capturing job-specific events at each client. Both user command events (e.g., job submission, abort commands) and critical learning events (e.g., round completion, aggregation steps) are recorded [48]. These audit logs contain event records with their timestamps, event type, and identity of the issuing party. This log structure means that, following any run, an institutional reviewer can verify which administrative commands were issued and by whom, which sites participated in each computation round, and that no raw data transfer commands were executed. Currently, the audit logs are accessible to clients as container logs, viewable via the Docker [31] or Singularity [32] interface. In a future release, these logs will be made available as part of the federated analysis workflow results returned through the NeuroFLAME interface.

#### Differential Privacy and Homomorphic Encryption (optional, not enabled by default)

NVFlare does not by default provide cryptographic protection against an authenticated participant that correctly executes the protocol but could in principle use the received parameter vectors to make inferences about client data. This is a known limitation of standard federated aggregation architectures, and NVFlare’s own documentation identifies it as a gap requiring additional cryptographic mechanisms in sensitive settings [47]. NVFlare provides a configurable data filtering pipeline operating at two points in every task exchange: task data filters, applied before data is passed to the local executor, and task result filters, applied to local outputs before transmission to the server [47].

**Differential Privacy (DP)** adds calibrated statistical noise to transmitted parameters before they leave a client site, providing a formal probabilistic bound on the amount of information about any individual that can be inferred from the transmitted values. The filter must be explicitly configured per job and is not activated in the current NeuroFLAME computation pipelines by default.

**Homomorphic Encryption (HE)** allows mathematical operations, including aggregation of model parameters, to be performed directly on encrypted data without decrypting it. NVFlare implements HE through integration with the TenSEAL library using the CKKS encryption scheme [47], designed for approximate arithmetic on floating-point vectors. Enabling HE requires generating TenSEAL context files and distributing public encryption keys to all participating sites during consortium provisioning. Integration of both DP and HE into NeuroFLAME’s computation pipelines is a defined priority for future development.

#### Unsafe Component Detection (optional, not enabled by default)

Unsafe component detection is available but requires explicit installation by each site. It operates through NVFlare’s event mechanism, allowing a custom component checker to inspect job components before they are built and raise an error if any are deemed unsafe [46]. Sites wishing to use this feature must define a checker class and register it in their site’s job resources.json configuration file.

Overall, NVFlare provides a practical and extensible federated learning framework that ensures baseline privacy through data localization and secure communication, while allowing users to augment its security model depending on the required threat assumptions.

### 4.7 Failure Handling and Operational Robustness

NeuroFLAME’s GUI includes an App Health check feature that reports each client’s critical component status prior to a run, such as Docker/Singularity connectivity and network connectivity. The Vaults admin console additionally reports each vault’s online/offline status. NVFlare performs pre-analysis checks before launching each sub-system to detect configuration errors before the workflow begins [7]. Computation developers can configure the minimum number of required clients per round according to the requirements of the computation. During execution, if a client site becomes unreachable or fails to respond, the NVFlare controller can be configured to continue with the remaining available sites rather than aborting the entire study. Jobs that fail can be re-queued by an authorized administrator via the FLARE admin console without re-provisioning the study. Provenance for each run including which sites participated in which rounds is preserved in the per-job workspace and returned to participants at completion, providing an auditable record of what executed and where.

**Fig. 2:**
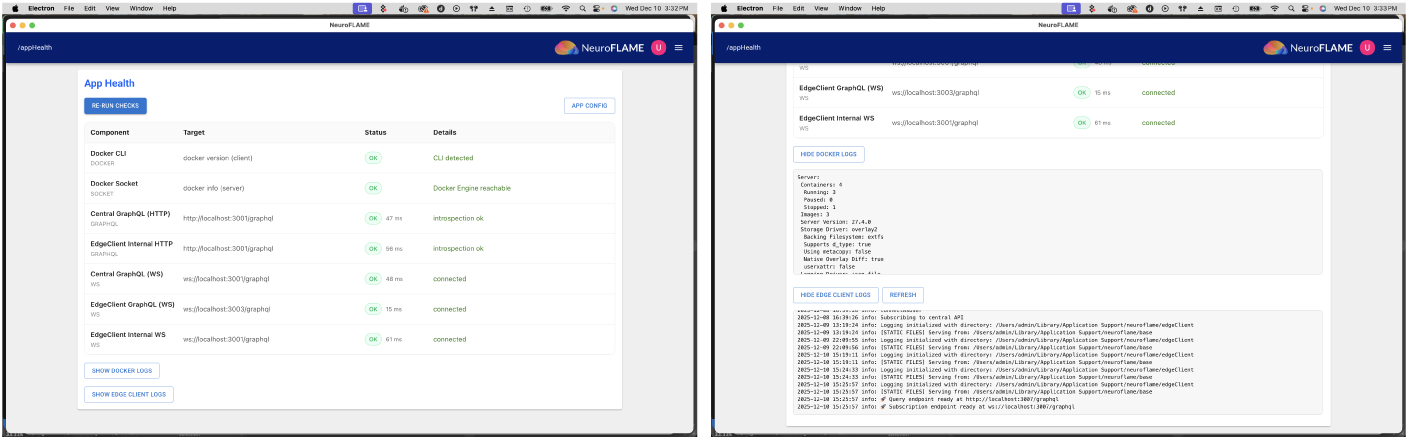
NeuroFLAME interface showing node health and real-time computation logs.

### 4.8 Workflow Execution Model

A federated computation in NeuroFLAME proceeds through a series of coordinated steps designed to maintain security, reproducibility, and operational robustness in collaborative environments. The standard execution flow is:

1. **Study Configuration**: Researchers utilize the desktop application to configure study parameters, select participating sites, and define the specific computation to be executed.
2. **Manifest Retrieval**: Edge Federated Clients at each participating site initiate outbound-only requests to the Central API to pull the job manifest and necessary runkits.
3. **Validation and Provisioning**: The Central Federated Client validates the study parameters and issues unique job manifests. The File Server automatically generates and distributes the required cryptographic startup kits to ensure secure communication.
4. **Container Orchestration**: Upon receiving the manifest, the Edge Federated Clients instantiate isolated containers using Docker (or Singularity for HPC environments). These containers encapsulate the entire computational workload and the NVFlare runtime environment.
5. **Secure Distributed Execution**: Once containers are active, they establish encrypted peer-to-peer channels using NVFlare secure communication protocols. Computations execute locally on the protected data at each site, with model updates or summary statistics exchanged according to the federated computation.
6. **Aggregation and Results**: The central server (or a designated lead site) aggregates the encrypted derivatives. Once the analysis reaches convergence or completion, the final results and full provenance tracking are returned to the participants via the desktop interface.

**Fig. 3:**
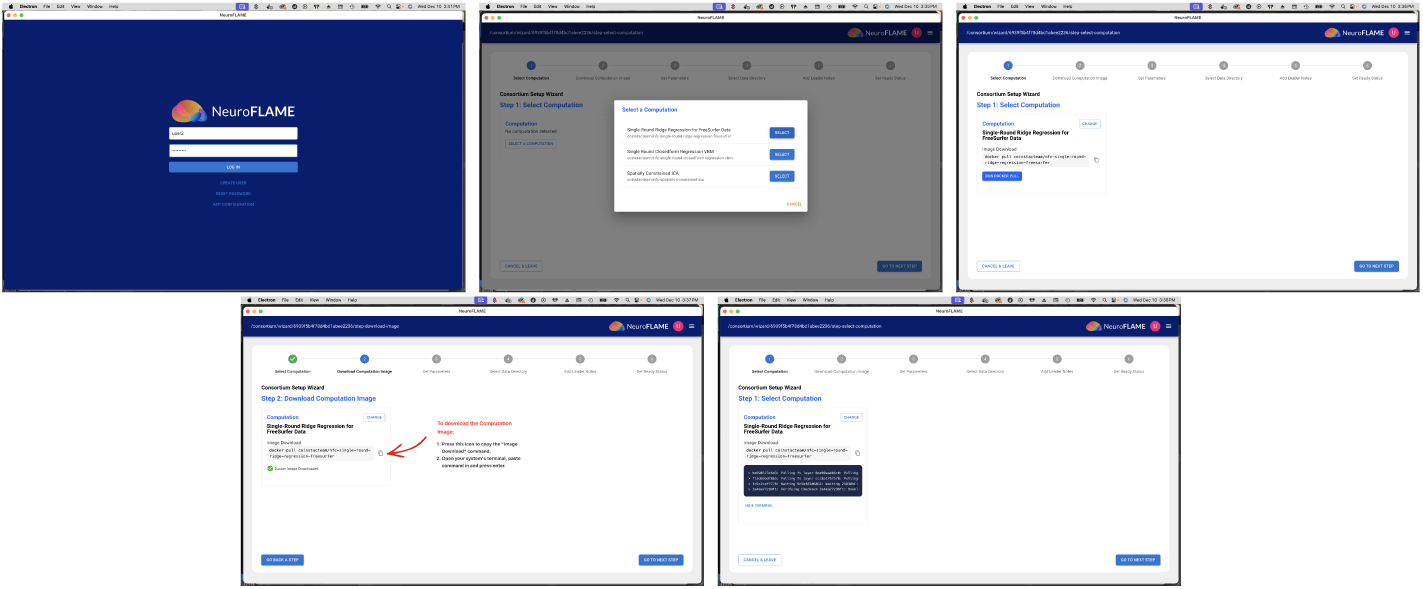
UI steps: Login, computation selection, and Docker image retrieval.

**Fig. 4:**
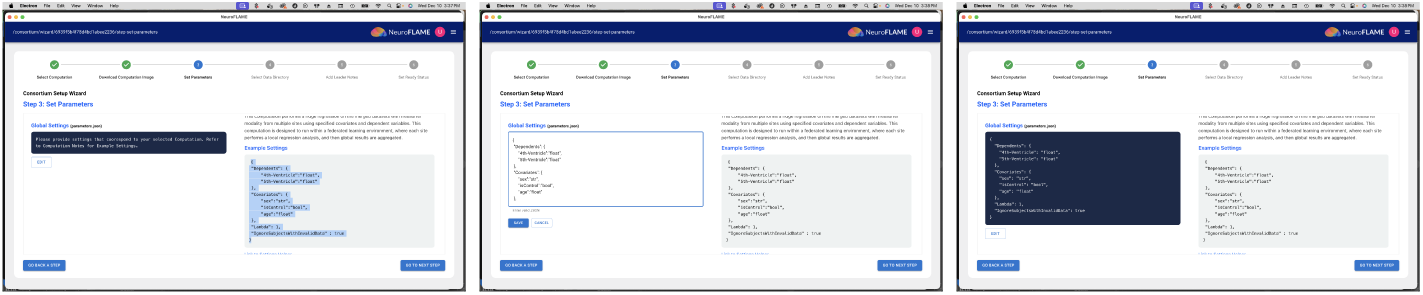
Parameter configuration.

**Fig. 5:**
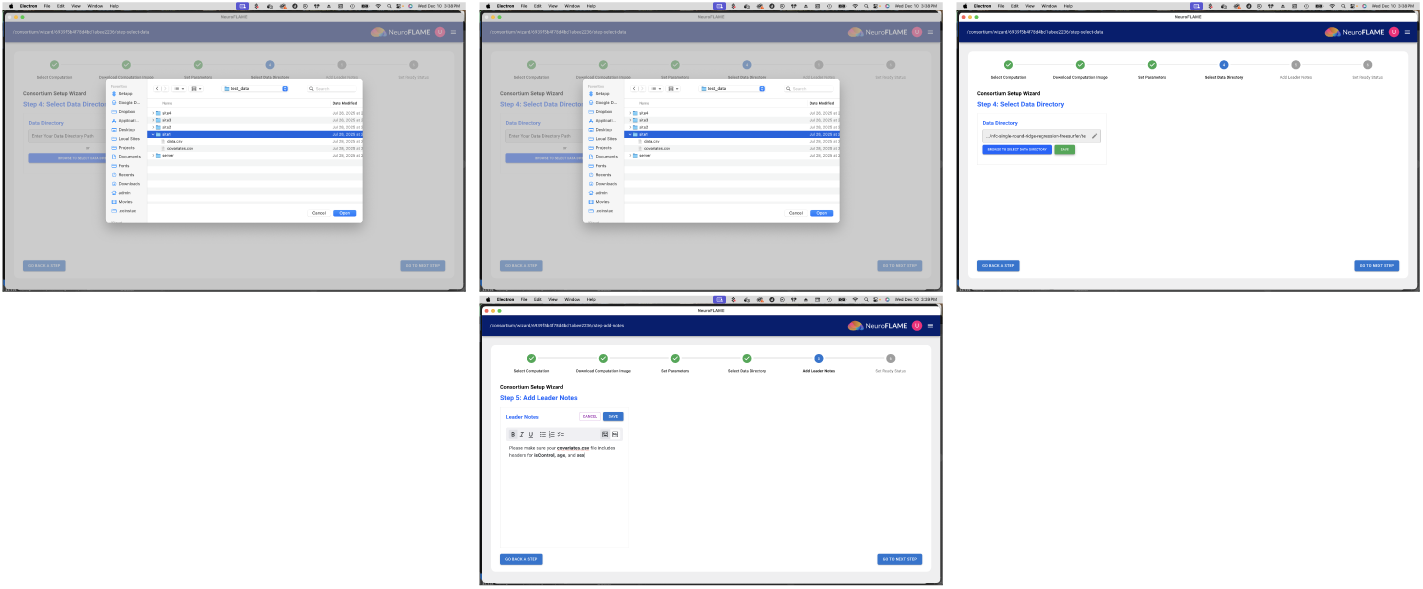
Local data directory selection.

**Fig. 6:**
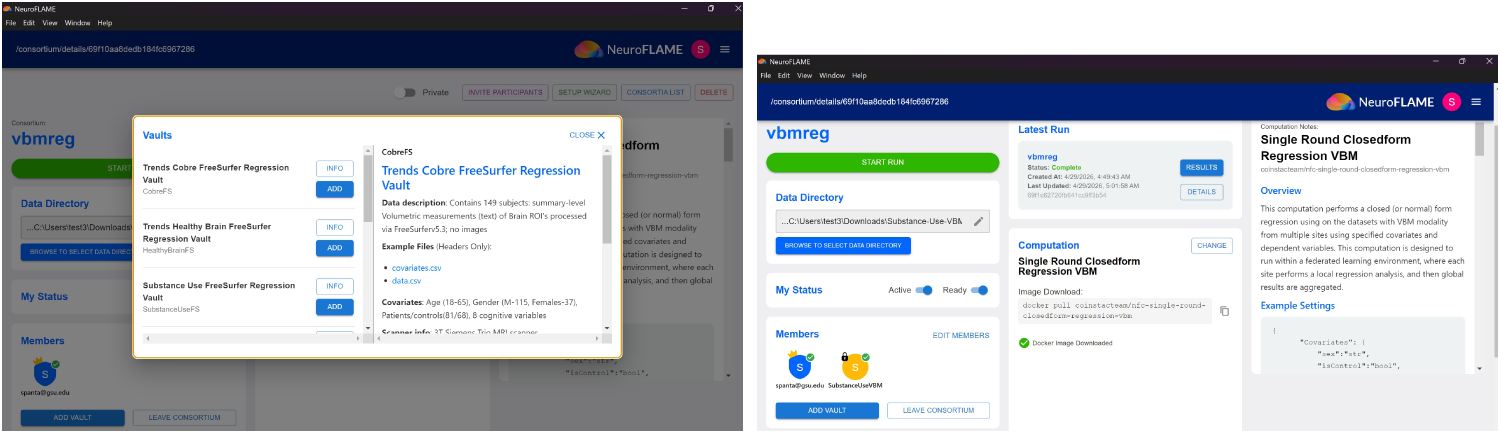
Add Vaults: browsing available vaults and viewing vault details within a consortium (left); consortium view after a vault has been added as a participating edge client (right).

**Fig. 7:**
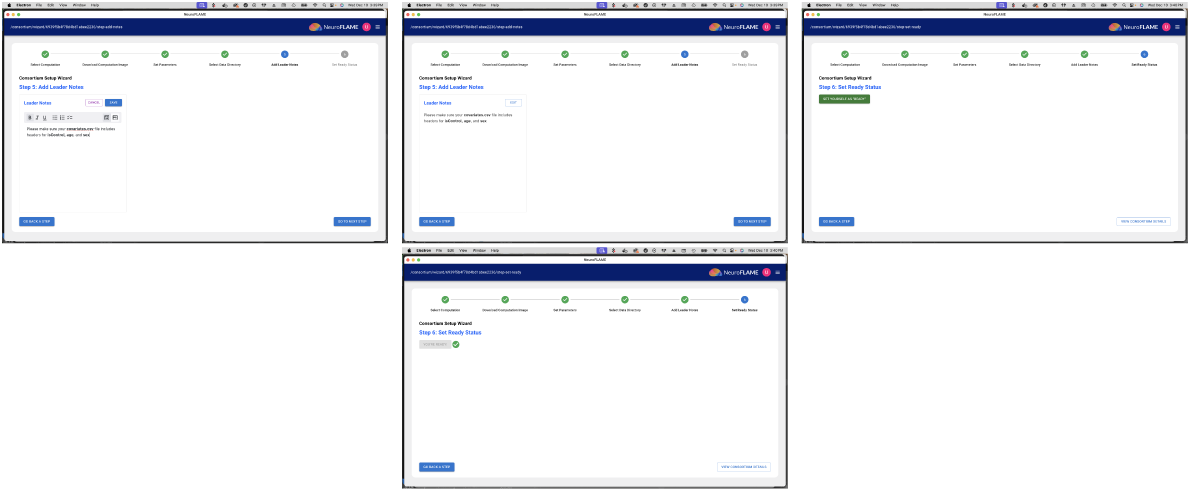
UI: add researcher notes and change to ready status.

**Fig. 8:**
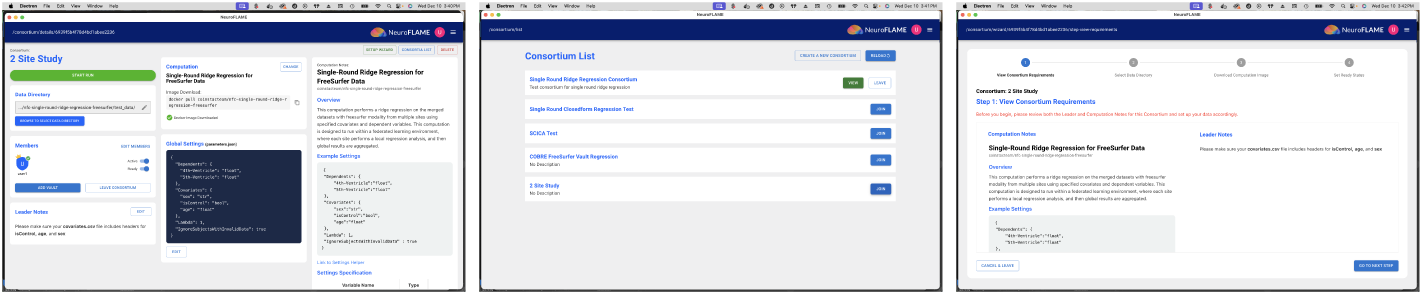
Finalizing consortium setup and viewing study details.

**Fig. 9:**
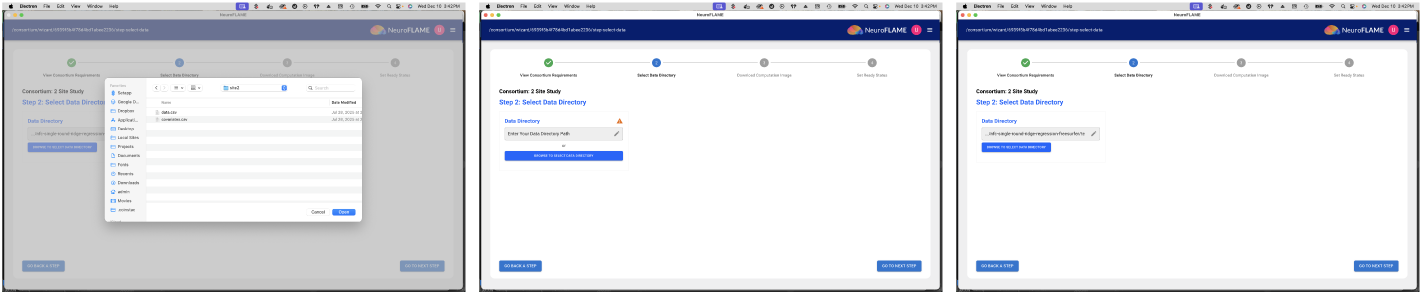
Consortium member workflow including requirement review and local data selection.

**Fig. 10:**
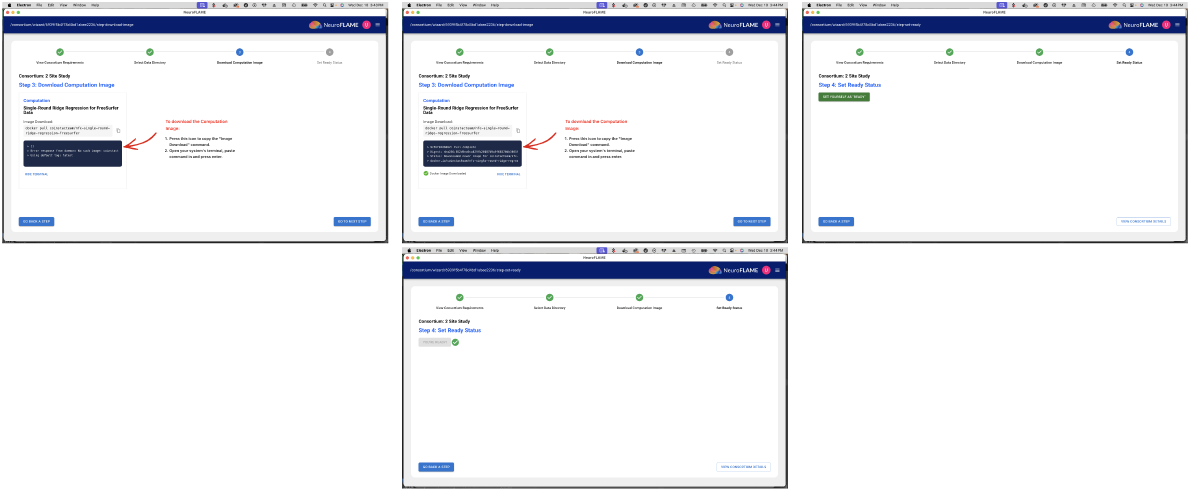
Member image download, status readiness, and active run monitoring.

## 5 Computation Development via NeuroFLAME

Our NeuroFLAME architecture is designed to lower the barrier of entry for researchers to implement and deploy federated neuroimaging analyses. By abstracting the complexities of peer-to-peer networking, certificate-based trust, and outbound-only communication at client sites, the platform allows developers to focus exclusively on the algorithmic logic of their study.

### 5.1 Simplified Integration via the Standardized Boilerplate

To facilitate the rapid development and integration of novel methodologies, NeuroFLAME introduces a standardized nvflare-boilerplate public repository (https://github.com/NeuroFlame/computation-nvflare-boilerplate) that can be used as a template for computation development. This repository serves as a containerized environment, pre-configured with the requisite environmental parameters and hierarchical directory structures necessary for NeuroFLAME runtime compatibility. The boiler-plate effectively decouples the “underlying infrastructure” from the “computation,” allowing an author to transition their local analysis to a federated context by modifying three primary functional modules:

- **Task Definition and Logic:** Implementation of site-specific local preprocessing or global aggregation steps within the application logic.
- **Data Handling:** Use of built-in data loaders to interact with local site datasets while maintaining privacy.
- **Job Configuration:** Leveraging automated configuration scripts (e.g., makeJob.py) to wrap the computation into a deployable container, which is then managed by the NeuroFLAME central orchestrator.

### 5.2 Architectural Advantages for New Authors

The NeuroFLAME architecture is uniquely beneficial for the scientific community for several reasons:

- **Infrastructure Abstraction:** Authors do not need to manage Docker orchestration, TLS certificate exchanges, or firewall configurations. The platform handles the “handshake” between participating sites and the central aggregator.
- **Native Neuroimaging Compatibility:** Unlike general-purpose federated learning frameworks, the boilerplate is pre-optimized for high-dimensional neuroimaging data (e.g., fMRI, sMRI), offering native support for common preprocessing pipelines and specialized neuroimaging file formats.
- **Standardized Validation:** A robust testing environment within the boilerplate (using local simulators) ensures that authors can validate their federated logic on synthetic or local data before deploying to a global multi-site deployment.

By adopting this boilerplate-driven approach, NeuroFLAME transforms the development of federated computations from a complex engineering task into a streamlined scientific workflow.

### 5.3 Existing Computations

NeuroFLAME currently supports a diverse library of containerized computations for federated neuroimaging analysis. Each computation is a decentralized algorithm packaged for execution across participating sites, enabling preprocessing, local statistical execution, controlled aggregation, harmonization, decomposition, prediction, or validation without centralizing raw files. NeuroFLAME analyses span multiple operational modes rather than a single class of federated method. Some computations use one-pass or low-communication aggregation, in which sites compute local summary quantities or sufficient statistics that are combined centrally. Others use multi-round iterative optimization, in which sites repeatedly compute local updates that are aggregated to fit a shared model, decomposition, or representation. We therefore use federated analysis as the umbrella term for the analytic scope of NeuroFLAME, computation to refer to a specific containerized decentralized algorithm, and federated learning for the subset of computations focused on shared model fitting or representation learning.

### 5.4 Federated Statistical Modeling

NeuroFLAME implements both single-round and multi-round federated regression models tailored for large-scale neuroimaging datasets.

- **FreeSurfer Ridge Regression (Single & Multi-Round)**: Designed for structural metrics (thickness, surface area, and volume). The single-round version averages local coefficients, while the multi-round version uses iterative gradient descent to converge on a global solution, offering high precision for distributed datasets.
- **Voxel-Based Morphometry (VBM) Regression**: A single-round, closed-form ordinary least squares (OLS) ridge regression. Local sites compute *X^T^ X* and *X^T^ Y* matrices from NIfTI files, which the controller aggregates to produce global beta maps, p-values, and *R*^2^ metrics.

### 5.5 Privacy-Enhanced Federated Model Fitting

The following suite of machine learning computations provides differentially private classification, automates label reclassification for diagnostic consistency, and facilitates multi-site data harmonization.

- **Differentially Private SVM (DP-SVM)**: For classification tasks involving highly sensitive data, NeuroFLAME provides a differentially private implementation of *L*^2^-regularized Logistic Regression and Support Vector Machines. This module uses objective or output perturbation to provide mathematical privacy guarantees. It trains local binary classifiers at each site and aggregates them at an “aggregator” site to form a global classifier without exposing subject-level features or labels.
- **Federated Label-Based Dimensional Prediction (Fed-LAMP)**: This is a federated extension of the label-noise filtering-based dimensional prediction (LAMP) framework [9] that enables multi-site analysis of functional network connectivity (FNC) data. The original LAMP method [9] identifies subjects exhibiting strong neuroimaging biomarkers consistent with their clinical labels, uses these “typical” subjects to construct a dimensional prediction model for mental disorders, and achieves enhanced classification accuracy by filtering label noise from neuroimaging data. Fed-LAMP adapts this pipeline to a privacy-preserving multi-site setting, enhancing the accuracy of typical-subject selection across institutions without centralizing raw neuroimaging data.
- **Federated ComBat (ComBat-DC / ComBatMega-DC)**: A federated implementation of the ComBat harmonization computation [49]. The module derives site-neutral means and variances to remove “site effects,” making data from different institutions statistically comparable while keeping raw datasets on-premises. Sites extract local summary statistics and share them with a central aggregator to compute site-neutral parameters, and the module then harmonizes the data locally.

### 5.6 Federated Decomposition and Multimodal Neuroimaging Fusion

The computations described below enable multivariate mapping of brain structure and function without pooling raw anatomical data. These modules focus on multivariate decomposition to identify functional and structural brain networks.

- **Decentralized Constrained Source-Based Morphometry (dcSBM)**: Performs a decentralized extension of constrained source-based morphometry. Each site conducts local constrained ICA using shared reference components derived from large external datasets, such as the UK Biobank. Statistical analyses identify significant group differences in structural network expression while maintaining near-identical findings to centralized approaches.
- **Spatially Constrained ICA (scICA):** Utilizes a joint optimization for independence and adherence to spatial templates (e.g., Neuromark [50]). This module bundles masking, variance removal, and static functional network connectivity (sFNC) computation into a managed pipeline.
- **Decentralized Constrained Joint ICA:** Implements a novel pipeline for multimodal fusion of functional and structural data, grounded in the NeuroMark Fusion framework [8]. It uses subject-space concentration to perform local constrained computations followed by federated averaging of the estimated unmixing matrices.

## 6 Results

### 6.1 VBM regression with Vault

In this section we will showcase federated analysis using the vault feature. We run a federated voxel-wise regression on pre-processed structural T1-weighted images with age, diagnosis, and gender as covariates. We will examine the relationship between grey matter volume with age and diagnosis.

This analysis utilizes a pre-curated, anonymized vault dataset (the TReNDS COBRE VBM Regression Vault) that is made available to NeuroFLAME users at all times. This vault is hosted on a private Amazon EC2 instance under the TReNDS Center organization and is ready for federated regression analysis. In this analysis, we have two users with different datasets located on their local machines, or nodes, as they will be referred to from here on. The images in these datasets are pre-processed through the VBM (Voxel-Based Morphometry) pipeline described in Section 6.1.2, so the datasets are ready for analysis at the point of access. A summary of the final datasets used in the analysis is outlined in the following section.

#### 6.1.1 Summary of datasets

##### TReNDS COBRE VBM Regression Vault

The vault hosts pre-processed grey matter images from 152 participants, approximately half healthy controls and half individuals diagnosed with schizophrenia, collected at the Mind Research Network as part of the Center of Biomedical Research Excellence (COBRE) study. [51]. In addition to imaging data, the vault includes demographic variables, symptom severity scales, and cognitive measures from which a user can build a design matrix; the present analysis is based only on age, sex, and diagnosis.

##### Johns Hopkins dataset

This dataset comprises 45 participants (27 healthy controls, 18 patients) scanned at Johns Hopkins University as part of a first-episode psychosis study. This is the final analytic sample, retaining only individuals diagnosed with schizophrenia (’SZ’) and healthy controls (’HC’) after pre-processing, quality control, and sub-selection to match the age, gender, and diagnosis distributions of the vault dataset. For this federated analysis, this dataset is hosted by a researcher in Albuquerque, NM, which serves as the second participating site.

##### Working memory study dataset

This dataset is drawn from the OpenNeuro collection *Working Memory in Healthy and Schizophrenic Individuals*, accession ds000115^1^ [52], collected at the Conte Center for the Neuroscience of Mental Disorders, Washington University in St. Louis [53]. Only the T1-weighted structural images were used. After pre-processing, quality control and matching with vault dataset distributions for for age,gender and diagnosis, 57 participants were retained (22 healthy controls, 35 patients). This dataset is hosted by a researcher at Emory University which serves as the third participating site.

#### 6.1.2 Single-round closed-form VBM regression

##### Pre-processing and quality control

All three datasets were prepared independently at their respective sites using the NeuroFLAME Preprocessor tool(Section 4.3) [41], and same pre-processing pipeline was applied at every site without any exchange of image data. Images were segmented into gray matter, white matter, and cerebrospinal fluid in SPM12 [42]. These images were spatially normalized to the standard Montreal Neurological Institute (MNI) template using a 12-parameter affine model, and smoothed with a 6 mm FWHM isotropic Gaussian kernel. For quality control, each individual’s smoothed normalized gray matter image was correlated with SPM12’s gray matter template, and the correlation value was computed. These values were examined for each cohort, and outliers were removed if they fell more than a certain number of standard deviations from the cohort mean.

##### Regression model

Voxel-wise associations between gray matter concentration and the covariates of interest were estimated using the single-round closed-form VBM regression computation in NeuroFLAME. The smoothed, normalized gray matter maps serve as input, with the gray matter value at each voxel as the dependent variable and age, gender, and diagnosis, together with an intercept, as independent variables. An absolute mask threshold of 0.2 was used to remove background noise, and input images were subsampled to 2 mm isotropic resolution to accelerate the analysis run time. These options are included in the computation to accommodate analysis with large datasets to speed up the run time.

The model is fit in a single communication round. Let *X_s_ ∈* R*^ns×p^* denote the design matrix at site *s* and *Y_s_ ∈* R*^ns×v^* the corresponding matrix of gray matter values over *v* voxels. Each site computes the cross-product matrices *X*^⊤^_*s*_*X_s_* and *X*^⊤^_*s*_*Y_s_* locally and transmits only these summary statistics to the aggregation server. Because both quantities are additive over sites, the server reconstructs the pooled normal equations exactly and solves the closed-form ordinary least squares estimator

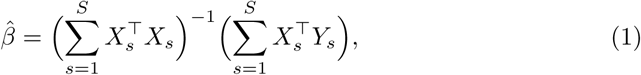

returning the pooled coefficients to the local nodes. No individual-level data leaves any site at any point, and the federated estimate is numerically identical to the result a pooled analysis on centralized data would produce. The design matrices were full rank at every site, so no regularization was required. Because the estimator is closed-form, a single round of communication is sufficient and no iterative optimization is involved.

##### Statistics returned

Voxel-wise *t* statistics are computed from the residual mean squared error and the design covariance, and converted to two-tailed *p* values under the *t* distribution. Results are reported as signed significance, *−* log_10_(*p*) *×* sign(*β*). The magnitude of a voxel encodes the strength of the statistical evidence and its sign carries the direction of the effect, so that positive and negative values indicate opposite directions of association. Model fit is summarized as *R*^2^ = 1*−* SSE*/*SST. Beta, signed significance, and *R*^2^ maps are written out as NIfTI volumes.

#### 6.1.3 Study configuration and execution

The federated analysis proceeds as follows.

1. Once a user creates a computation, they add the TReNDS COBRE VBM Regression Vault and select the computation, in this case, single-round closed-form VBM regression. The parameters and options for the computation are supplied in JSON format through the GUI. This allows the model specification, covariates, mask threshold, and subsampling factor to be changed easily.
2. The other member is then added to the consortium, and each add the local path to their dataset, in this case the Johns Hopkins data and the working memory study data respectively. This data is restricted to the local machines and never leaves the site. Only the derivative summary statistics outlined in Section 6.1.2 are shared between sites and central NeuroFLAME server.
3. Once the leader runs the computations, the regression analysis runs locally at each site via Docker or Singularity containers, depending on how the user’s machine is setup. When the run finishes, each site receives a copy of its own local statistics along with the global summary statistics, in this case voxel-wise beta values for each covariate in the regression model, together with the corresponding signed significance and *R*^2^ maps. These are provided in nifti format and .png files. The results of this federated analysis are shown in the following section.

#### 6.1.4 Site-level results

Each site receives its own local statistics in addition to the global result. Figure 11 shows the signed significance maps computed on the TReNDS COBRE VBM Regression Vault with 152 individuals. The gray matter volume decreases with age, especially in the basal ganglia[54], and patients show reduced volume relative to controls, consistent with previous reports in this cohort and in the wider literature [55, 56].

**Fig. 11:**
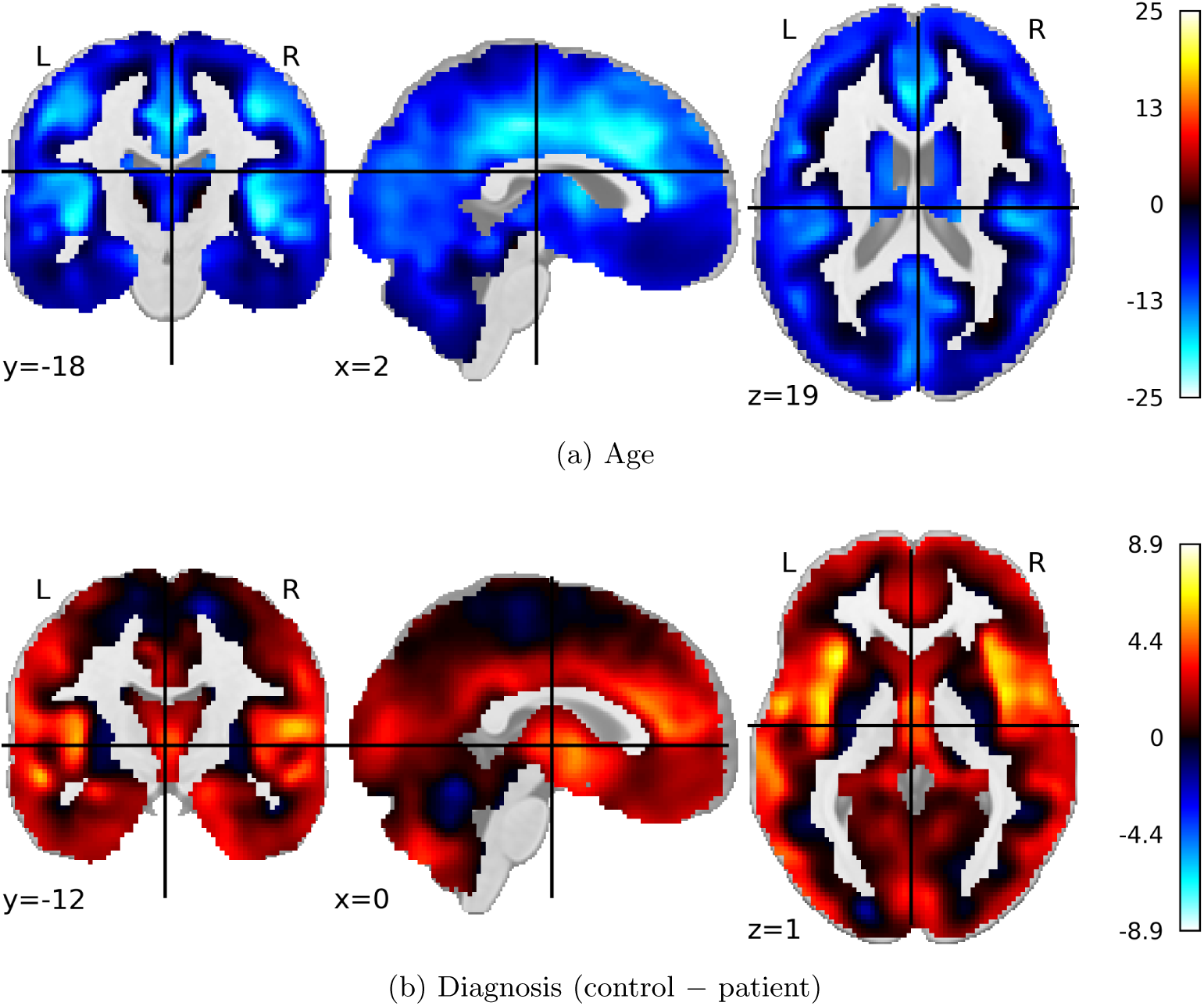
**TReNDS COBRE VBM Regression Vault (***n* = 152). Signed significance maps, *−* log_10_(*p*) *×* sign(*β*), for the **(a)** age and **(b)** diagnosis covariates. The magnitude of a voxel reflects the strength of the statistical evidence and its sign gives the direction of the effect. For age, negative values (blue) indicate that gray matter volume decreases with increasing age. For diagnosis, positive values (red) indicate that control gray matter volume exceeds patient gray matter volume, and negative values (blue) the reverse.

Figure 12 shows the corresponding maps for the Johns Hopkins dataset. The direction of both effects is reproduced, with negative age associations across gray matter volume,, and reduced gray matter volume in patients and reduced patient volume; the evidence at any individual voxel is weaker and more spatially scattered than in the vault, as expected with smaller dataset of 45 individuals over a narrower age range.

**Fig. 12:**
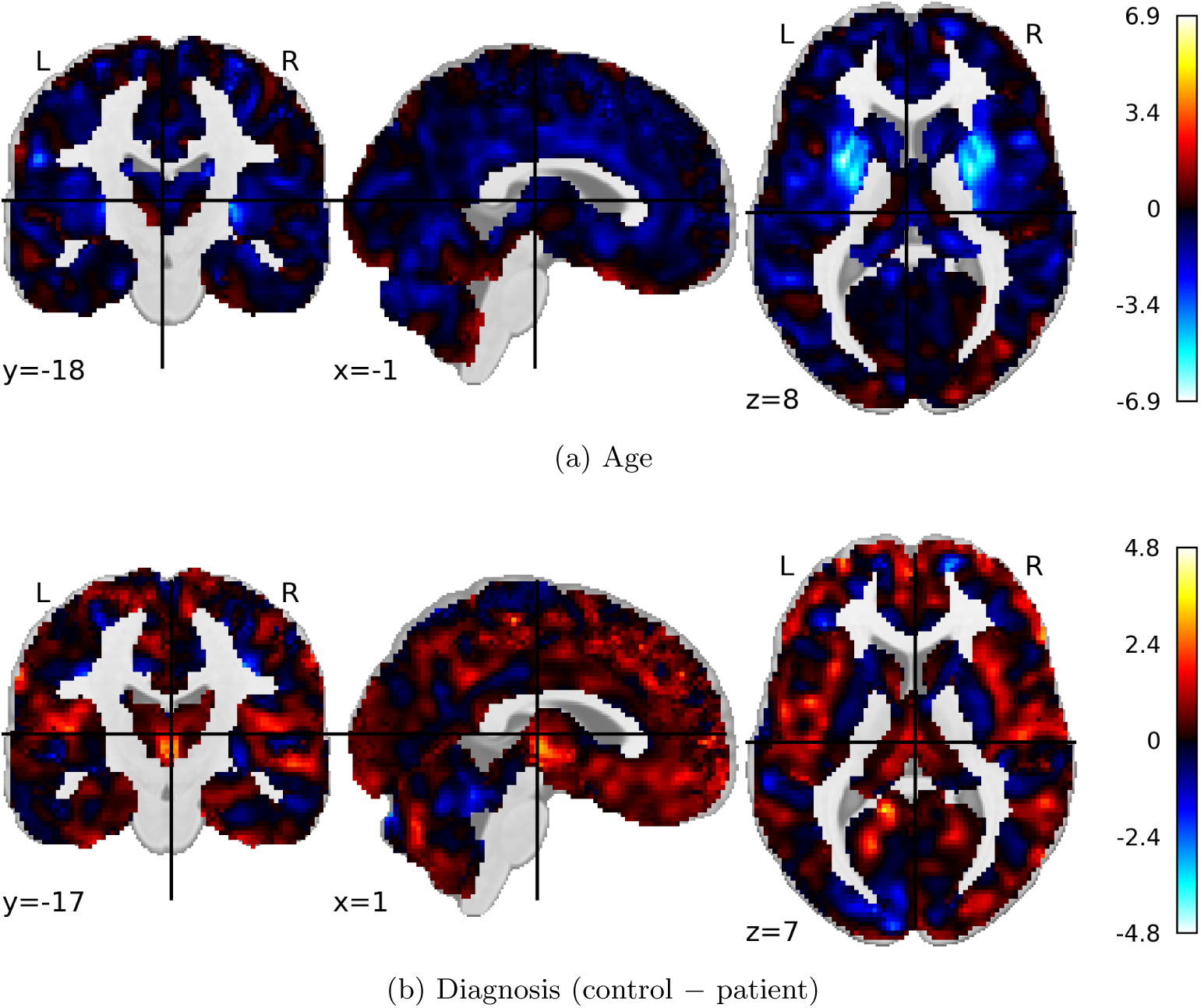
**Johns Hopkins dataset (***n* = 45). Signed significance maps for the **(a)** age and **(b)** diagnosis. Sign conventions and colour scale interpretation are as in Figure 11; note the reduced colourbar range relative to the vault.

Figure 13 shows the maps for the working memory study dataset with 57 individuals. The same pattern is recovered with correspondingly weaker voxel-wise evidence.

**Fig. 13:**
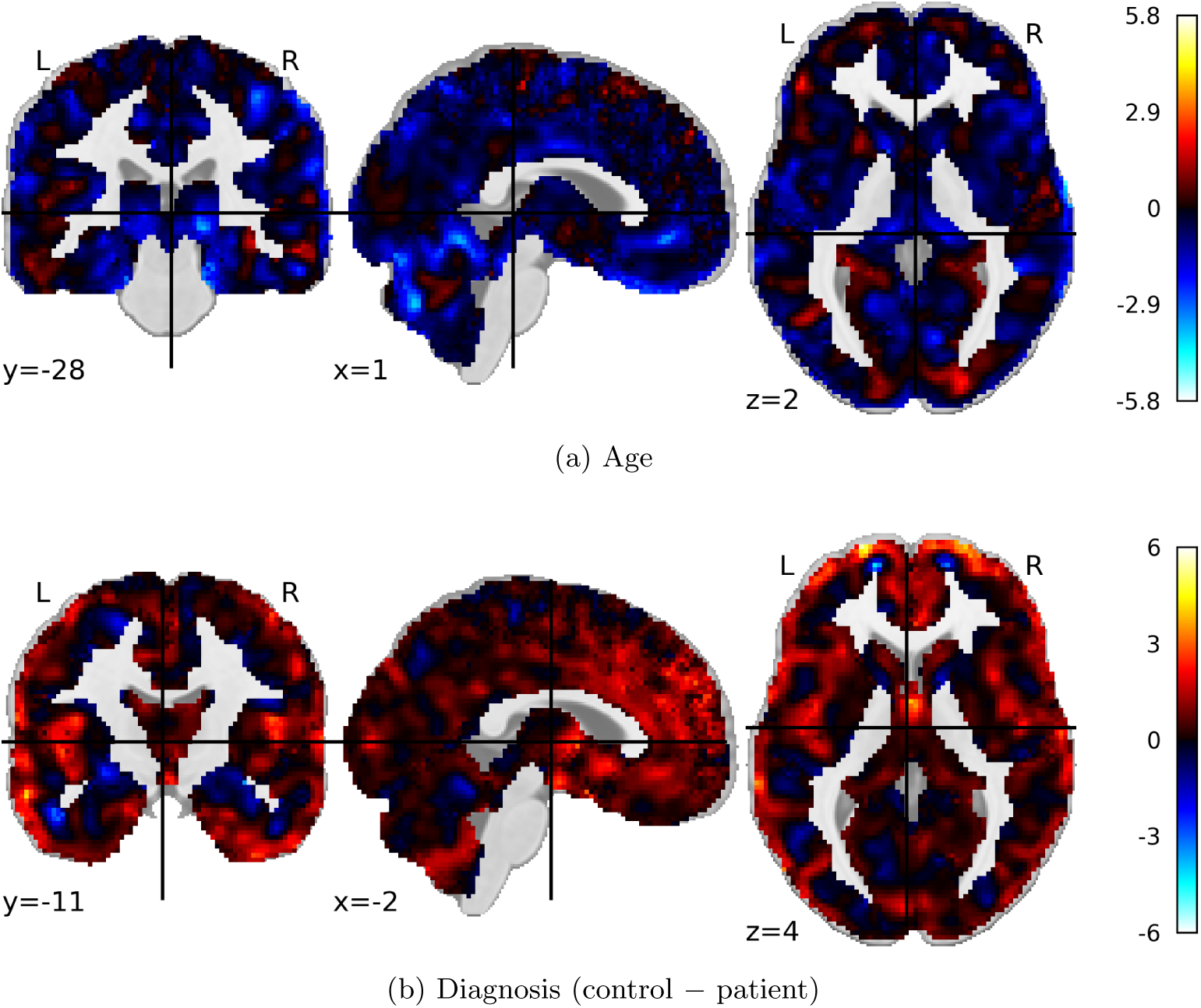
**Working memory study dataset (**ds000115, *n* = 57). Signed significance maps for the **(a)** age and **(b)** diagnosis. Sign conventions and colour scale interpretation are as in Figure 11.

#### 6.1.5 Federated global statistics

Combining the vault with both local datasets in a single federated regression raises the sample from 152 to 254 participants, and the patient group from 83 to 136 (Table 1). The global statistics returned to each site are shown in Figure 14.

**Table 1:** Summary of the final datasets entering the federated regression. Age is given as mean *±* SD. HC, healthy controls; SZ, patients.

| Dataset | Node | N | HC | SZ | Age (years) | M | F |
| --- | --- | --- | --- | --- | --- | --- | --- |
| TReNDS COBRE VBM Regression Vault | Vault | 152 | 69 | 83 | $38.3 \pm 12.5$ | 115 | 37 |
| Johns Hopkins | Local | 45 | 27 | 18 | $25.5 \pm 4.3$ | 31 | 14 |
| Working memory (ds000115) | Local | 57 | 22 | 35 | $24.4 \pm 3.6$ | 46 | 11 |
| **Federated total** | | **254** | **118** | **136** | $32.5 \pm 11.8$ | **204** | **67** |

**Fig. 14:**
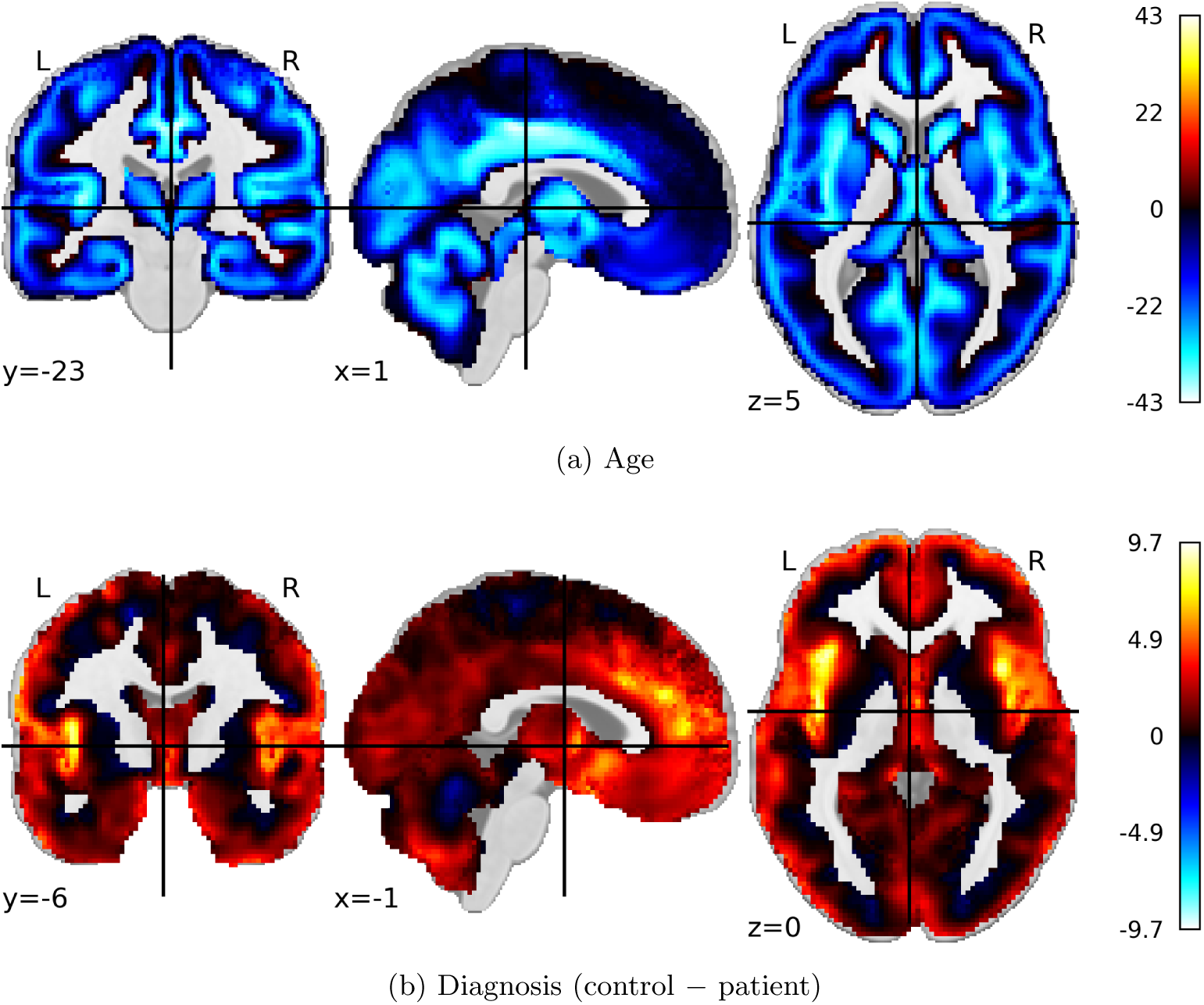
Federated global statistics. Signed significance maps for the **(a)** age and **(b)** diagnosis from the combined analysis of the TReNDS COBRE VBM Regression Vault, the Johns Hopkins dataset, and the working memory study dataset (*n* = 254). Sign conventions are as in Figure 11. Note the expanded colourbar range relative to the site-level maps.

The federated result is consistent in direction and anatomical distribution with the site-level analyses, and the statistical evidence is substantially stronger: for both the age and diagnosis covariates, the signed significance is markedly greater than at any contributing site alone. The diagnosis maps show a spatially distinct pattern in the insular temporal and frontal regions[57]. This result demonstrates the statistical utility of federated analysis for questions that require larger samples than any single site can provide on its own.

### 6.2 Decentralized Constrained Joint ICA

We implement a novel pipeline for decentralized multi-modal fusion of functional and structural imaging data. Following the example of decentralized neuromark ICA, we utilize a template generated from a large sample (*N >* 5000) of non-affected individuals from the UK Biobank dataset [8]. The NeuroFLAME implementation thus consists of two separate stages:

#### 6.2.1 Compute sFNC and GMV Derivatives

For rsfMRI, spatially constrained ICA was applied using the GIFT toolbox [58] and the NeuroMark 1.0 template [50] to extract subject-level spatial maps for each of the 53 intrinsic connectivity networks (ICNs) defined in the template, along with their associated activation time courses. FNC was computed as the pairwise Pearson correlation of ICN time courses. For sMRI, T1-weighted images were preprocessed with SPM12 [59]; modulated gray matter probabilistic segmentation maps were extracted and smoothed with a Gaussian kernel (FWHM = 12 mm) to obtain GMV maps. Subjects with excessive head motion (greater than 3 degrees rotation or 3 mm translation, or mean framewise displacement exceeding 0.3 mm) were excluded.

#### 6.2.2 Utilize Decentralized Constrained ICA to Estimate a Decentralized Joint Fusion

The NeuroFLAME implementation of this pipeline operates in two coordinated stages. In the first stage, each participating site independently computes sFNC derivatives from rsfMRI data and GMV derivatives from sMRI data using the preprocessing pipelines described above, without transferring any raw imaging files. In the second stage, the decentralized guided joint ICA computation is executed: each site performs local constrained ICA using the shared NeuroMark Fusion template as a spatial prior, estimating local unmixing matrices that map subject-level multimodal features onto the template components. These locally estimated unmixing matrices are then transmitted to the central aggregator, which combines them via federated averaging to compute a global estimate of the joint unmixing solution [8]. This approach extends the NeuroMark template-guided, reproducible network estimation into the multimodal domain, and makes the comparison of fusion results across federated sites tractable by anchoring all local solutions to the same normative template.

#### 6.2.3 Results

Figure 15 summarizes the correspondence between the decentralized constrained joint ICA results and a pooled (centralized) reference, evaluated across varying numbers of sites (2, 4, 8, 16, and 32). Across all site configurations, the federated implementation achieves high spatial correlation with the pooled result, with correlations exceeding 0.95 for up to 16 sites and above 0.90 even at 32 sites. Additionally, we apply the template in a decentralized fashion to the A4/Learn data set[60] and measured associations of the estimated GMV and sFNC components with age. The loadings for one joint component was highly associated with age, and the associated loading distribution, GMV and sFNC maps are provided in figure 16.

**Fig. 15:**
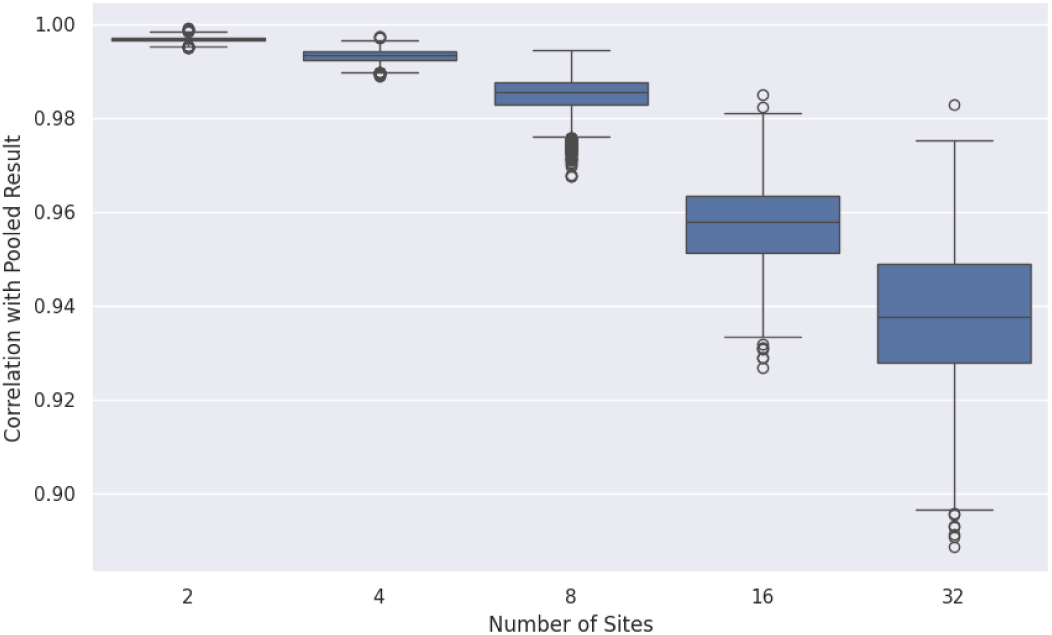
Joint ICA Federated vs. Pooled results: correlation with pooled reference across numbers of simulated sites (2, 4, 8, 16, 32).

**Fig. 16:**
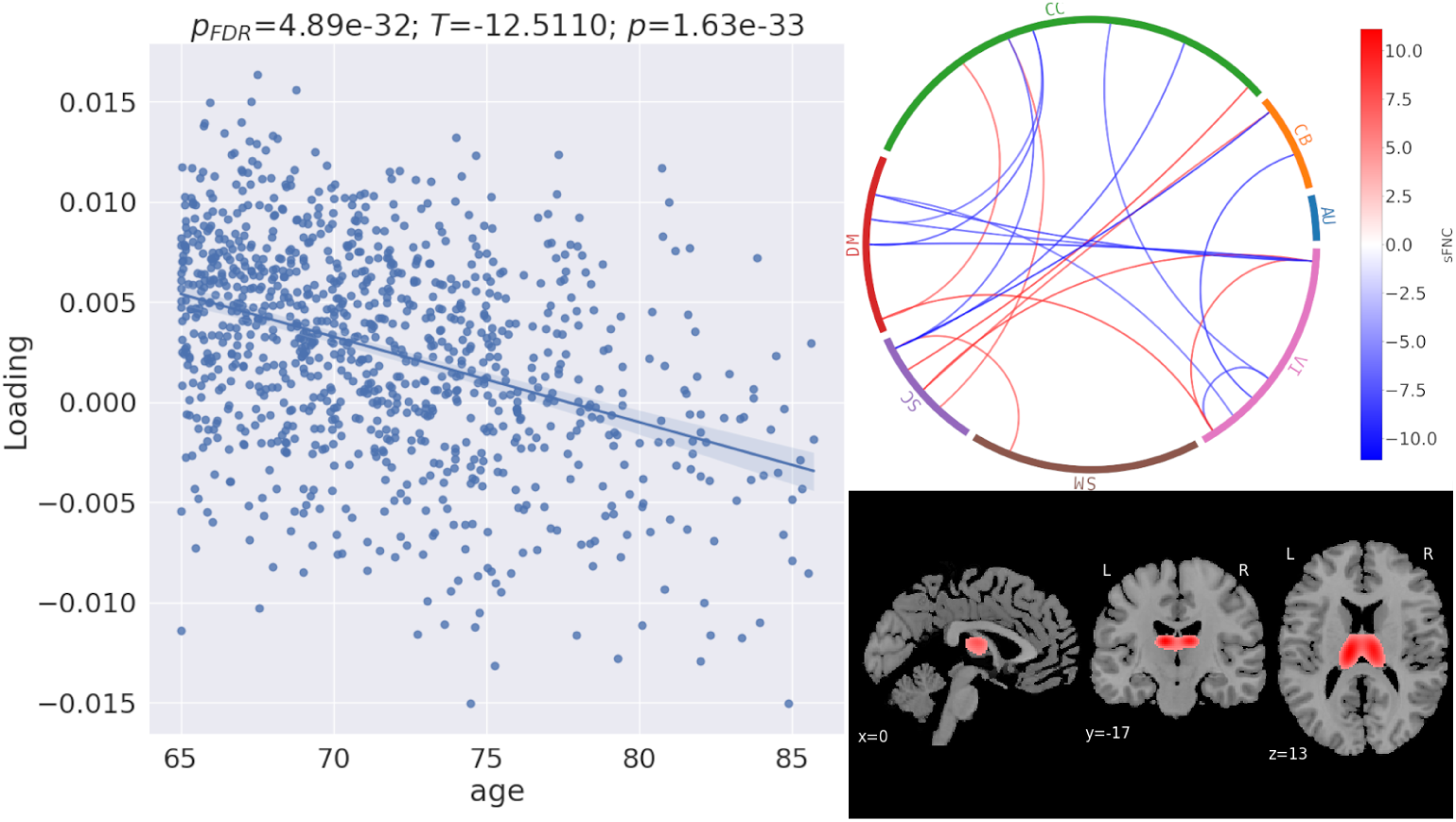
An example of the joint components and associated loadings for decentralized constrained joint ICA in static functional network connectivity and gray matter, which were significantly associated with age in the A4/Learn data set.

These results demonstrate that the decentralized guided joint ICA pipeline in NeuroFLAME’s framework preserves the scientific behavior of the centralized approach across a range of realistic multi-site configurations.

### 6.3 Label-Based Dimensional Prediction (Fed-LAMP)

Federated LAMP (Fed-LAMP) extends the original label-noise filtering-based dimensional prediction framework [9] into a privacy-preserving multi-site setting for neuroimaging studies. The core idea is to preserve the logic of LAMP [9] while avoiding centralized sharing of raw subject-level fMRI data, which is often restricted. In the original LAMP workflow [9], resting-state fMRI is transformed into Functional Network Connectivity (FNC) features, and a complete random forest (CRF)-based label-noise filtering strategy is used to identify typical subjects whose imaging patterns are more consistent with their clinical labels. These typical subjects’ within-group averages are used to form representative centroids for each class. In our federated reformulation, these same operations are performed locally at each site, building local model summaries without moving raw neuroimaging data. Only compact centroids are shared with the aggregator, allowing the global workflow to remain privacy compliant. Each subject is compared with the centroids of healthy and patient-derived typical subjects, and the relative similarity to these reference groups is converted into a dimensional score. For a new dataset or an unseen subject, the score provides a principled way to estimate how closely that subject resembles a healthy or disease-related neuroimaging pattern. Adaptive thresholding is applied: instead of using a fixed cutoff, the aggregated score distribution is examined to derive a global threshold that clearly separates healthy, patient-like, and boundary subjects. This thresholding step is crucial because it prevents overconfident relabeling of borderline cases.

The results show that the federated implementation preserves the essential behavior and utility of the centralized LAMP method [9] while eliminating the need for raw data pooling. A two-sample *t*-test (Figure 17) of FNC features yielded sharper, more consistent group-difference maps after relabeling, indicating greater intra-group compactness and improved inter-group separation. Comparisons between centralized and federated relabeling (Figure 18) further showed that the resulting FNC patterns were nearly identical, demonstrating that the federated design preserves the scientific behavior of the original method rather than approximating it loosely. Overall, this project shows that LAMP can be successfully decentralized, enabling robust, privacy-preserving neuroimaging analysis across institutions without compromising the original methodological intent.

**Fig. 17:**
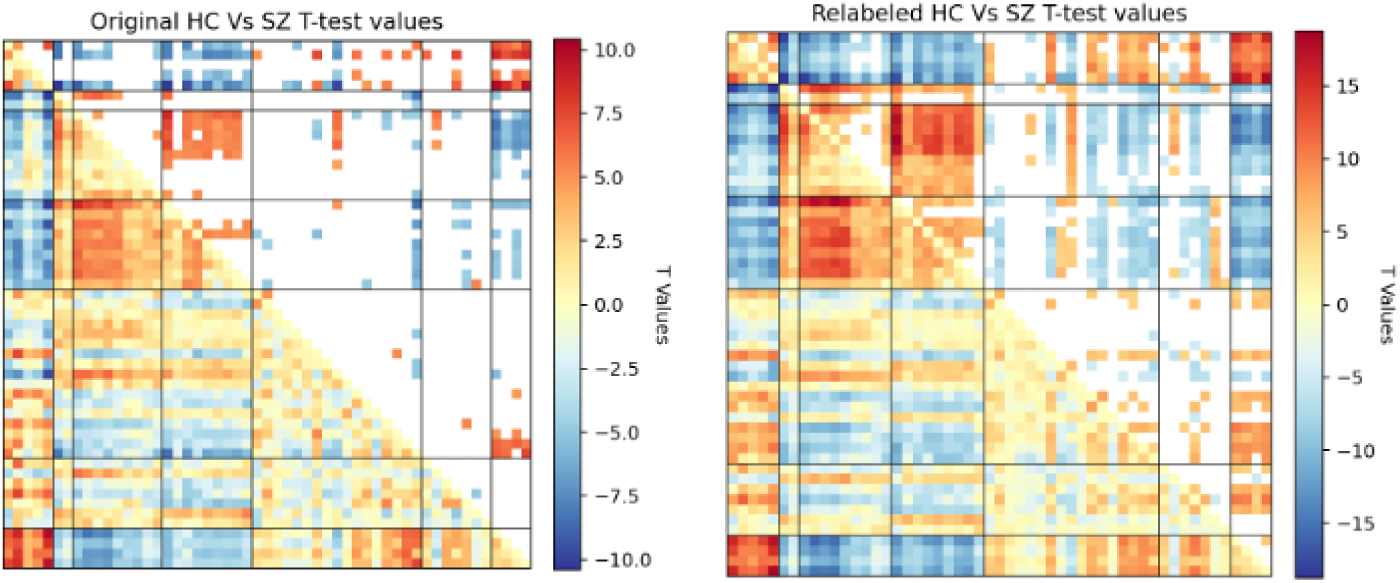
*t*-test heatmaps demonstrating group separability improvements using Fed-LAMP [9].

**Fig. 18:**
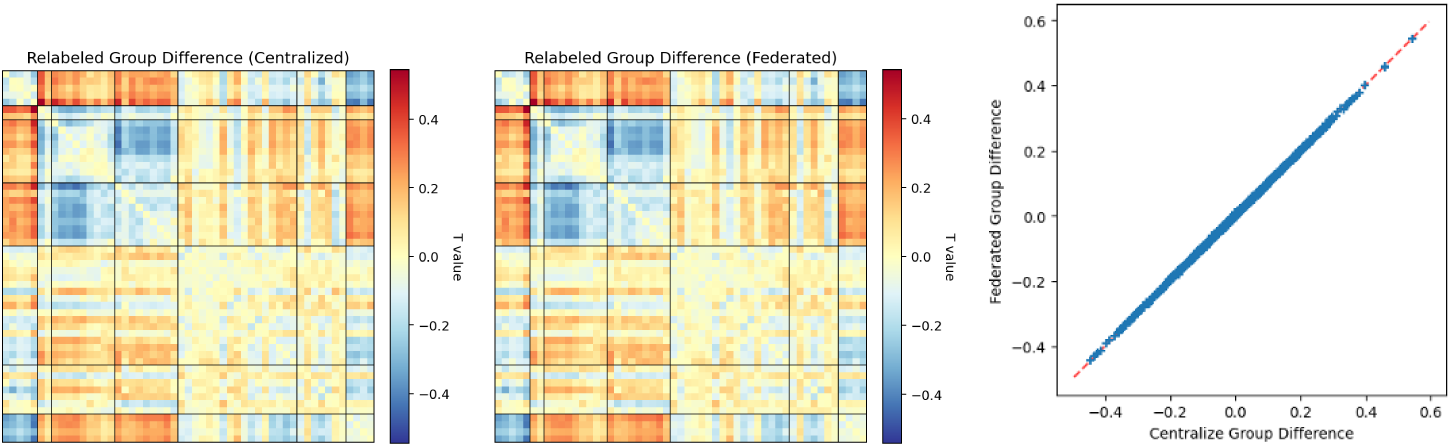
Comparison of Centralized and Federated LAMP results [9].

## 7 Conclusion

The development of NeuroFLAME addresses a critical implementation gap in federated neuroimaging computation, where many proposed approaches remain confined to technical proofs of concept, simulated deployments, or workflows that are difficult to operate under real-world institutional governance and clinical IT constraints. By bridging the divide between enterprise-grade security and user-centric design, our platform provides a sustainable model for production-ready, collaborative neuroimaging computation and analysis. At the system level, the integration of the NVFlare framework ensures a secure, extensible privacy-preserving communication backbone [46]. Simultaneously, the neuroscience-focused GUI and containerized computation model lower the barriers to entry for domain experts, allowing researchers to prioritize statistical insights over complex communication logistics. The empirical validation through diverse federated computations supported by NeuroFLAME demonstrates that it maintains high scientific consistency with established centralized approaches [8, 9]. Ultimately, NeuroFLAME provides the infrastructure to realize the potential of privacy-preserving federated computation for large-scale, multi-institution collaborative neuroimaging and health research, while remaining compatible with both federated analysis and federated learning workflows.

## Data and Code Availability Statement

The TReNDS COBRE VBM Regression Vault is hosted by the TReNDS Center and is available to NeuroFLAME users for federated analysis; the underlying COBRE data are described in Aine et al. [51].

The Johns Hopkins first-episode psychosis data are not publicly available.

The working memory study data are openly available from OpenNeuro under accession ds000115 (https://openneuro.org/datasets/ds000115) [52].This data was obtained from the OpenfMRI database. Its accession number is ds000115. Users of this dataset are additionally asked to cite the associated publication [53] and the repository [61, 62].

The NeuroFLAME preprocessor tool as outlined in Section 4.3 is used to preprocess all structural MRI datasets in this work. This tool is openly available at https://github.com/NeuroFlame/neuroflame-preprocessor; the analyses reported here used release 0.1.2 [41].

The data used in the decentralized constrained joint ICA analysis (Section 6.2) were provided by the A4/LEARN Study. These data are not publicly available and were used under a data use agreement. Qualified researchers may apply for access through the A4 Study data portal at https://www.a4studydata.org.

Two datasets were analyzed in the federated LAMP analysis: COBRE and FBIRN. The COBRE data are described in Aine et al. [51], and the FBIRN data are drawn from the Function Biomedical Informatics Research Network data repository [63]. The scripts referred to in the manuscript are available on GitHub in the following repositories: https://github.com/NeuroFlame/nfc-label-noise-filtering. The NeuroFLAME platform code is available at https://github.com/NeuroFlame/.

## Footnotes

1 https://openneuro.org/datasets/ds000115

